# LRP2 expression in melanoma is associated with a transitory cell state, increased T cell infiltration, and is upregulated by IFNγ signaling

**DOI:** 10.1101/2025.02.21.639443

**Authors:** Martin Q. Rasmussen, Marie L. Bønnelykke-Behrndtz, Camilla Merrild, Ida Tvilling, Julie N. Christensen, Morten M. Nielsen, Jeanette B. Georgsen, Nina Naumann, Johann M. Gudbergsson, Anders Etzerodt, Jakob S. Pedersen, Russell W. Jenkins, Søren E. Degn, Søren K. Moestrup, Henrik Schmidt, Torben Steiniche, Mette Madsen

**Author notes:** Corresponding author. All communication to.

## Abstract

Low density lipoprotein receptor-related protein 2 (LRP2) is a 600 kilodalton multi-ligand endocytic membrane receptor expressed in several cell types during fetal development, including neuroepithelial cells, and in select absorptive epithelial cells in the adult. In epithelial cancers, LRP2 expression is associated with a differentiated tumor cell state and better prognosis. In previous work, we found that while LRP2 is not expressed in benign naevi, it is frequently acquired in melanoma. However, the molecular drivers of LRP2 expression in melanoma and characteristics of LRP2-expressing melanoma have yet to be described. Here, we show that LRP2 expression is related to a transitory melanoma cell state defined by co-expression of melanocyte lineage and neural crest transcriptional programs. Further, we reveal that melanoma LRP2 expression is increased in T cell-inflamed tumors, and is directly upregulated through interferon-gamma signaling. Correlation of melanoma LRP2 expression with clinicopathological variables demonstrates that LRP2 expression is associated with low Breslow thickness and low clinical stage in primary melanomas. Taken together, the present study describes the characteristics of LRP2-expressing melanoma and reveals interferon gamma signaling as a novel strong positive regulator of LRP2 expression in melanoma.

**Significance:** Melanoma cells often acquire LRP2 expression but the drivers of LRP2 expression in this setting and characteristics of LRP2-expressing melanoma remain unclear. Here, we show that LRP2 expression is related to a transitory melanoma differentiation cell state. Further, LRP2 expression in melanoma correlates with a T cell-inflamed tumor microenvironment and LRP2 expression in melanoma cells can be directly increased by interferon-gamma. In addition, LRP2 expression is associated with less advanced histopathological characteristics of melanoma. These findings encourage future studies on LRP2 in settings with increased interferon signaling, in particular in melanoma metastases following immunotherapy.

## Introduction

Melanoma cells can adopt transcriptional cell states that resemble the embryonic origin of melanocytes from the neural crest (Tsoi et al., 2018). Melanoma dedifferentiation is defined by loss of melanocyte-lineage transcriptional programs, including genes involved in pigment production, which are controlled by microphthalmia-associated transcription factor (MITF), the master regulator of melanocyte development (Levy et al., 2006). Melanoma cells also transiently re-activate a neural crest-like transcriptional program through the course of dedifferentiation. Four melanoma cell state classifications based on the activity of the melanocyte-lineage and neural crest-like gene programs have been established: melanocytic (melanocyte-lineage^high^, neural crest-like^low^), transitory (melanocyte-lineage^high^, neural crest-like^high^), neural crest-like (melanocyte-lineage^low^, neural crest-like^high^) and undifferentiated (melanocyte-lineage^low^, neural crest-like^low^) (Tsoi et al., 2018).

Both tumor intrinsic (e.g. complex genetic backgrounds) and tumor extrinsic factors (e.g. inflammatory signals, hypoxia, and nutrient availability) contribute to the transcriptional heterogeneity of melanoma cells (Rambow et al., 2019). Melanoma dedifferentiation can be driven by inflammatory cytokine signaling, including tumor necrosis factor (TNF) and interferon-gamma (IFNγ), and is observed following immune checkpoint blockade (Grasso et al., 2021; Kim et al., 2021) and adoptive T cell transfer in preclinical models (Landsberg et al., 2012) and patients (Mehta et al., 2018).

Low-density lipoprotein receptor-related protein 2 (LRP2), also known as megalin, is a 600 kilodalton multi-ligand endocytic membrane receptor (Nielsen et al., 2016) expressed in absorptive epithelial cells in the adult. In epithelial cancers, LRP2 is highly expressed in differentiated subtypes, while epigenetic silencing related to methylation of a conserved CpG site in the first intron of the LRP2 gene is observed in dedifferentiated subtypes (Rasmussen et al., 2023). In the context of melanoma, we previously showed that LRP2 is frequently expressed in melanoma cells and tumors, while it is rarely expressed in benign naevi (Andersen et al., 2015). In the melanocytic lineage, LRP2 is expressed in neuroepithelial cells, which give rise to the neural crest, at an early stage of melanocyte differentiation during fetal development (Willnow et al., 1996). Still, the characteristics of LRP2-expressing melanoma and mechanisms driving LRP2 expression in melanoma remain unclear.

In the present study, we set out to characterize LRP2-expressing melanoma. Based on transcriptional analysis of melanoma cell lines and tumors, we find that high LRP2 expression in melanoma is related to a transitory transcriptional cell state defined by co-expression of melanocyte-lineage and neural crest-like gene programs. Interestingly, we detect a positive correlation between LRP2 expression in melanoma and the abundance of tumor-infiltrating lymphocytes and activation of inflammatory signaling pathways within the tumor microenvironment. Cell culture cytokine stimulation experiments demonstrate that IFNγ can directly increase and sustain LRP2 expression in melanoma cells. Finally, immunohistochemical analysis of primary melanomas reveals that LRP2 expression in melanoma cells is associated with low Breslow thickness and low clinical stage. Taken together, this study describes the molecular and clinicopathological features of LRP2-expressing melanomas and reveals IFNγ as a novel strong positive regulator of LRP2 expression in melanoma.

## Results

### LRP2 expression is associated with a transitory melanoma cell state

LRP2 expression is associated with a differentiated cell state in epithelial cancers (Rasmussen et al., 2023), but the relationship between LRP2 expression and melanoma differentiation cell state has not been examined. To address this, we analyzed bulk RNA sequencing data from a panel of melanoma cell lines (n = 53) representing various transcriptional melanoma cell states (Tsoi et al., 2018) (**Fig. 1A**). *LRP2* was infrequently expressed in the most differentiated melanocytic melanoma cell lines, consistent with lack of *LRP2* expression in benign naevi (Andersen et al., 2015). High *LRP2* expression was observed in a subset of transitory melanoma cells, which co-express high levels of melanocyte lineage and neural crest-related gene programs. *LRP2* was neither expressed in neural crest-like or undifferentiated melanoma cells (**Fig. 1B**).

**Figure 1.**
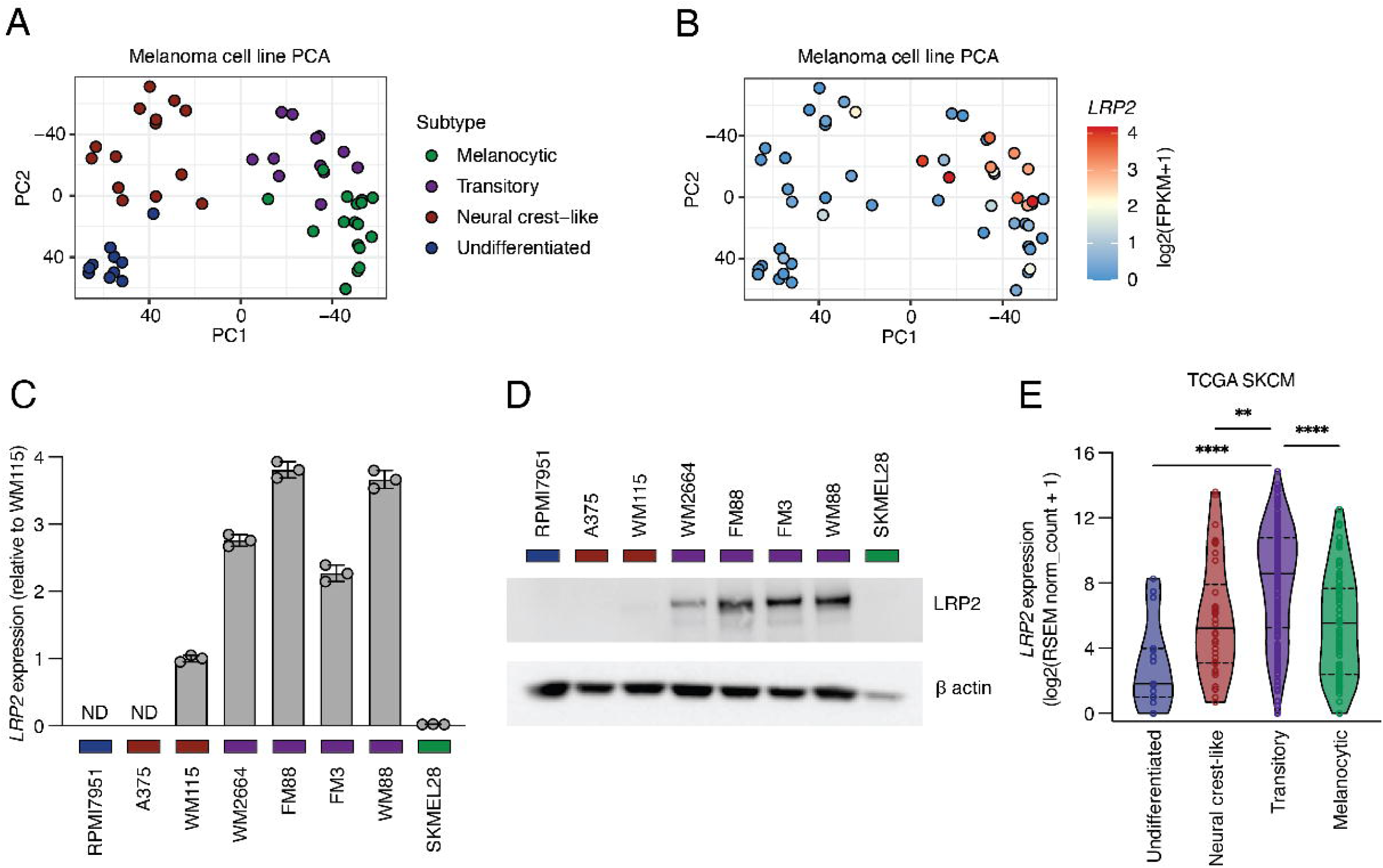
LRP2 expression is associated with a transitory melanoma cell state. **A.** PCA plots of bulk RNA sequencing data from melanoma cell lines color coded for transcriptional cell state classifications (melanocytic, transitory, neural crest-like or undifferentiated). **B.** PCA plot as in A color coded for *LRP2* expression (log2(FPKM+1). **C.** *LRP2* expression across eight melanoma cell lines determined by RT-qPCR. RQ values are normalized to WM115. Mean (bars) ± SEM (error bars) and individual values (circles) are shown (n = 3 technical replicates). **D.** Whole-cell lysate immunoblots for LRP2 (upper panel) and β actin loading control (lower panel) across eight melanoma cell lines. **E.** Violin plot of *LRP2* expression (log2(RSEM norm_count+1) in TCGA SKCM classified by transcriptional cell state (melanocytic, transitory, neural crest-like or undifferentiated)(one-way ANOVA followed by Dunnett’s multiple comparisons test). **P < 0.01, ****P < 0.0001. ND = Not detected. Color coding: green = melanocytic, purple = transitory, red = neural crest-like, blue = undifferentiated.

To validate LRP2 expression in transitory melanoma cells, we assembled a panel of melanoma cell lines representing melanocytic (SKMEL28), transitory (WM88, FM3, FM88, and WM2664), neural crest-like (WM115 and A375) and undifferentiated (RPMI7951) cell states (**Supplementary Fig. 1A**). High *LRP2* transcript levels were detected in transitory melanoma cell lines (WM88, FM3, FM88, and WM2664) and modest levels in one of two neural crest-like melanoma cell lines (WM115) (**Fig. 1C**). LRP2 protein levels were assessed by whole-cell lysate immunoblotting using a mouse monoclonal anti-human LRP2 antibody. Consistent with the *LRP2* transcript levels, LRP2 protein expression was observed in transitory melanoma cell lines (WM88, FM3, FM88 and WM2664), as well as minimal expression in the neural crest-like cell line WM115, but no detectable expression in other melanoma cell lines (**Fig. 1D**).

Next, we assessed the relationship between *LRP2* expression and melanoma differentiation cell state in human metastatic melanoma tumors in The Cancer Genome Atlas Skin Cutaneous Melanoma (TCGA SKCM) cohort. High *LRP2* expression was observed in transitory tumors compared to melanocytic, neural crest-like, and undifferentiated tumors (**Fig. 1E**). Taken together, these observations indicate that LRP2 expression is associated with a transitory cell state in melanoma.

### LRP2-expressing melanomas have increased immune infiltration

To further characterize LRP2-expressing melanomas, we performed gene set enrichment analysis of *LRP2*-correlated genes in TCGA SKCM (**Supplementary Table 1**). The top enriched pathways were related to immune responses and immune signaling, such as adaptive immune response, lymphocyte-mediated immunity, and IFNγ response (**Fig. 2A-B**), indicating a more inflamed tumor microenvironment in *LRP2*-expressing melanoma. To define the immune infiltrate in *LRP2*-expressing tumors, we used microenvironment cell population counter (MCPcounter), which uses the expression of cell type-specific marker genes to provide a relative score for major immune cell subsets in bulk RNA sequencing samples (Becht et al., 2016). We dichotomized TCGA SKCM into *LRP2*-high and *LRP2*-low groups using an upper quartile cutoff and compared cell population abundance. *LRP2*-high melanomas were generally enriched in monocyte, myeloid, and lymphoid cell populations with the greatest enrichment in cytotoxic lymphocytes (**Fig. 2C, Supplementary Fig. 2A-J**). We also applied a previously defined T cell-inflammation signature for the classification of TCGA SKCM tumors (Spranger et al., 2015) and observed higher *LRP2* expression in T cell-high compared to T cell-low melanomas (**Fig. 2D**).

**Figure 2.**
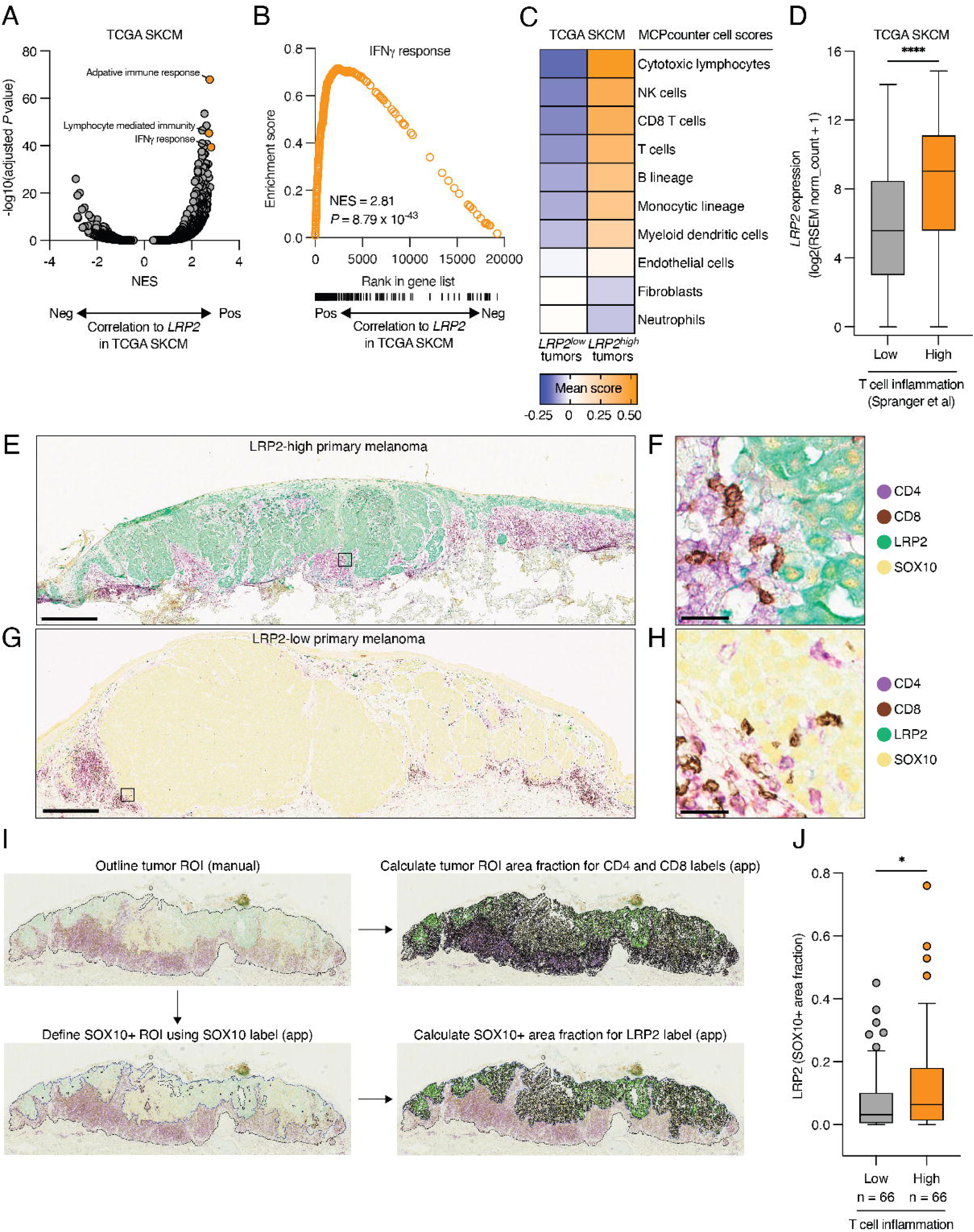
LRP2-expressing melanomas have increased immune infiltration. **A.** Volcano plot of gene set enrichment analysis results for *LRP2* correlated genes in TCGA SKCM. NES = Normalized Enrichment Score. Gene sets of interest are highlighted in orange. **B.** Mountain plot of the IFNγ response gene set for *LRP2* correlated genes in TCGA SKCM. NES = Normalized Enrichment Score. **C.** Heatmap of mean absolute MCPcounter scores for *LRP2*-low and *LRP2*-high tumors (upper quartile cutoff) in TCGA SKCM. **D.** Boxplot of *LRP2* expression (log2(RSEM norm_count+1) in T cell inflammation-high and T cell inflammation-low groups in TCGA SKCM (unpaired t-test). **E-H.** Image of four-color multiplex immunohistochemistry analysis for CD4 (purple), CD8 (DAB), LRP2 (teal) and SOX10 (yellow) in primary melanomas. **E.** Image of a LRP2-high primary melanoma. Box indicates zoomed area displayed in panel F. Scale bar = 500 μm. **F.** Zoom of the image in panel E. Scale bar = 25 μm. **G.** Image of a LRP2-low primary melanoma. Scale bar = 500 μm. Box indicates zoomed area displayed in panel H. **H.** Zoom of the image in panel G. Scale bar = 25 μm. **I.** Schematic of digital image analysis workflow for multiplex immunohistochemistry analysis of primary melanomas stained for SOX10 (yellow), LRP2 (teal), CD4 (purple) and CD8 (DAB). **J.** Boxplot of LRP2 (SOX10+ area fraction) in T cell inflammation-high and T cell inflammation-low groups from multiplex immunohistochemistry analysis of primary melanomas (Mann-Whitney test). The boxplot lines represent the lower quartile, median, and upper quartile. Whiskers extend to 1.5 times above or below the interquartile range. *P < 0.05, ****P < 0.0001.

To validate the positive correlation between *LRP2* gene expression and T cell inflammation in melanoma, we performed a four-color multiplex immunohistochemistry analysis of SOX10, LRP2, CD4, and CD8 in sections of formalin-fixed paraffin-embedded (FFPE) primary melanomas (**Fig. 2E-H**) using a cohort of 150 patients diagnosed with primary melanoma (**Supplementary Table 2**). One section of each tumor was scored for CD4 and CD8 T cells (fraction of bulk tumor region) and LRP2 (fraction of SOX10+ tumor region) (**Fig. 2I**). The abundance of CD4 and CD8 T cells showed a strong positive correlation (**Supplementary Fig. 2K**), consistent with previous characterizations of the immune infiltrate in primary melanomas (Nsengimana et al., 2018). Based on this, we stratified primary melanomas into T cell-high and T cell-low and compared LRP2 expression in SOX10+ melanoma cells. LRP2 expression in SOX10+ melanoma cells was higher in T cell-high versus T cell-low tumors (**Fig. 2J**), consistent with observations in the TCGA SKCM cohort. In combination, these results demonstrate increased expression of LRP2 in T cell-inflamed melanomas.

### IFN**γ** increases LRP2 expression in melanoma cells

T cell-secreted inflammatory cytokines, including IFNγ, can modulate the differentiation state of melanoma cells (Kim et al., 2021; Landsberg et al., 2012; Mehta et al., 2018; Tsoi et al., 2018). Based on the positive correlation between T cell infiltration, IFNγ signaling, and LRP2 expression in melanoma, we hypothesized that IFNγ can directly upregulate LRP2 expression in melanoma.

To test this hypothesis, we exposed a panel of melanoma cell lines to IFNγ and measured the expression of melanoma differentiation state marker genes (*MITF*, *MLANA*, *NGFR,* and *AXL*) and *LRP2*. As expected, IFNγ signaling could drive melanoma cell dedifferentiation, as indicated by downregulation of the melanocyte lineage markers *MITF* and *MLANA* and upregulation of the neural crest/dedifferentiation markers *NGFR* and *AXL* (Supplementary Fig. 3A-D). IFNγ increased *LRP2* expression in six of the seven cell lines tested (**Fig. 3A**). We observed increased *LRP2* expression following IFNγ-treatment in melanoma cell lines with intact JAK1/2 (**Supplementary Fig. 3E**), but not in melanoma cell lines with endogenous or engineered mutations in JAK1/2 (**Supplementary Fig. 3F**), in line with the canonical JAK-STAT signaling pathway being the principal mediator of IFNγ-driven gene expression changes (Grasso et al., 2021). By immunoblotting we confirmed increased LRP2 protein levels upon IFNγ-stimulation in WM2664 (**Fig. 3B**) and FM88 (**Fig. 3C**) melanoma cells.

**Figure 3.**
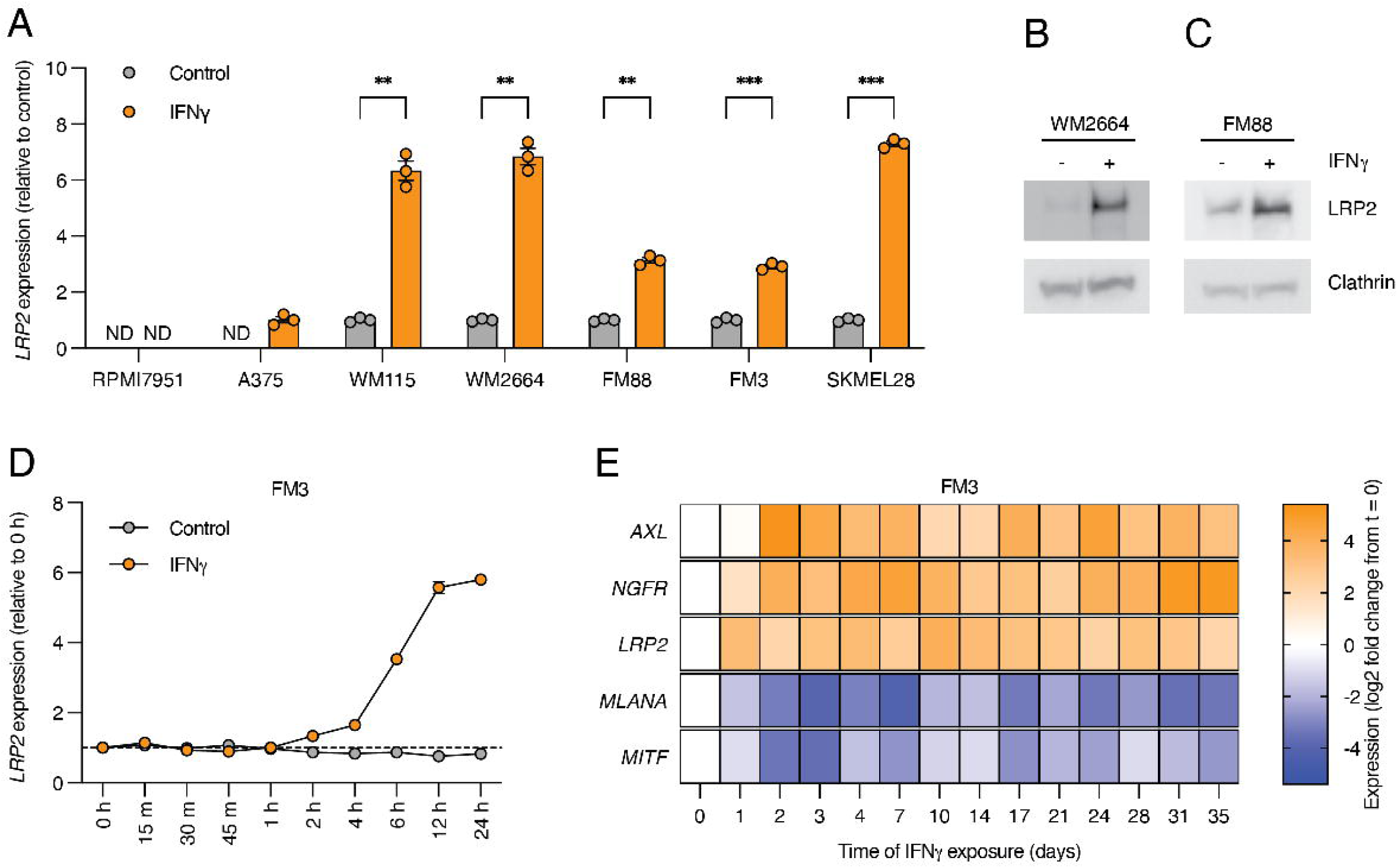
IFN**γ** increases LRP2 expression in melanoma cells. **A.** Bar plot of *LRP2* expression (relative quantities) determined by RT-qPCR in the indicated melanoma cell lines treated with vehicle control or IFNγ for 48 hours (multiple unpaired t-tests). Mean (bars) ± SEM (error bars) and individual values (circles) are shown (n = 3 technical replicates). *LRP2* relative quantities for A375 in the IFNy treated condition are set to 1 as LRP2 was not detected in the vehicle control. **B-C.** Whole-cell lysate immunoblots of LRP2 (upper panel) and clathrin loading control (lower panel) for the indicated melanoma cell lines treated with vehicle control or IFNγ for 48 hours. **D.** *LRP2* expression (relative quantities) determined by RT-qPCR during a 24-hour time course in cells treated with vehicle control or IFNγ. Mean (circles) ± SEM (error bars) are shown (n = 3 technical replicates). No error bars are displayed when the error is smaller than the size of the bar. **E.** Heatmap of gene expression (log2 fold change from t = 0) determined by RT-qPCR during a 5-week time course (n = 3 technical replicates). **P < 0.01, ***P < 0.001. ND = not detected.

We subsequently examined IFNγ-mediated *LRP2* induction in melanoma cells in a 24-hour time-course experiment. *LRP2* levels began to increase after two hours and continued to increase until the end of the experiment at 24 hours (**Fig. 3D, Supplementary Fig. 3G**). In an extended time-course experiment, continuous IFNγ-treatment sustained increased *LRP2* expression in melanoma cells until the final time point at five weeks (**Fig. 3E**). Collectively, these results demonstrate that IFNγ increases and sustains elevated LRP2 expression levels in melanoma cells.

### LRP2 is associated with low Breslow thickness and low tumor stage in primary melanoma

Finally, we applied the results from our four-color multiplex immunohistochemistry analysis of primary melanomas to assess the relation between melanoma cell LRP2 expression (LRP2 area fraction of SOX10+ tumor region) and clinicopathological variables (**Supplementary Table 2)**. We evaluated the association between LRP2 and established prognostic factors in primary melanoma, specifically Breslow thickness and ulceration. Thin melanomas (Breslow thickness < 1 mm) exhibited higher LRP2 expression compared to those with higher Breslow thickness (> 2 mm) (**Fig. 4A**). LRP2 expression was not associated to ulceration (**Fig. 4B**). We also assessed the association between LRP2 expression and clinical stage, observing higher LRP2 expression in stage I-melanomas compared to stage II-or stage III-melanomas (**Fig. 4C**). LRP2 was not associated to patient age, sex, anatomical location or histologic type (**Supplementary Fig. 4A-D**).

**Figure 4.**
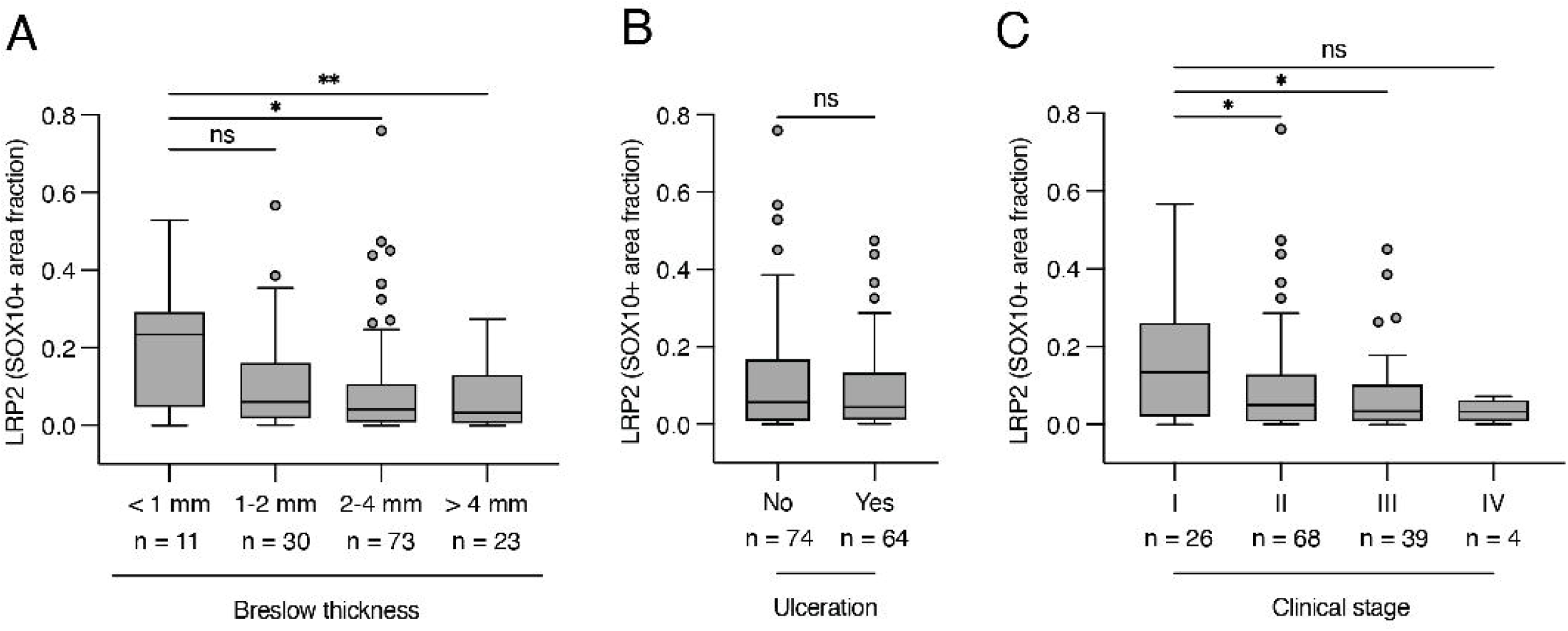
LRP2 is associated with low Breslow thickness and low clinical stage in primary melanoma. **A.** Boxplot of LRP2 (SOX10+ area fraction) expression by Breslow thickness categories in primary melanoma (one-way ANOVA followed by Dunnett’s multiple comparisons test). **B.** Boxplot of LRP2 (SOX10+ area fraction) by ulceration status in primary melanoma (Mann-Whitney test). **C.** Boxplot of LRP2 (SOX10+ area fraction) expression by clinical stage in primary melanoma (one-way ANOVA followed by Dunnett’s multiple comparisons test). The boxplot lines represent the lower quartile, median, and upper quartile. Whiskers extend to 1.5 times above or below the interquartile range. ns = not significant, *P < 0.05, **P < 0.01. ns = not significant.

## Discussion

Here, we show that LRP2 expression in melanoma is related to a transitory transcriptional cell state defined by concurrent expression of melanocyte lineage and neural crest-like gene expression programs. LRP2 is expressed at an early stage in the melanocyte lineage in neuroepithelial cells in the embryo (Willnow et al., 1996). Therefore, LRP2 expression in transitory melanoma cells might reflect re-activation of an embryonic neuroepithelial/neural crest transcriptional program involving LRP2.

We also found that melanoma LRP2 expression is correlated with T cell infiltration and IFNγ signaling in the tumor microenvironment and can be directly upregulated and maintained by continuous IFNγ stimulation. Previous work has demonstrated that T cell recognition of a minor fraction of tumor cells and IFNγ secretion to the tumor microenvironment has long-range effects resulting in IFNγ-sensing by a large fraction of the tumor (Hoekstra et al., 2020). In line with this concept, T cell activation following immunotherapy and secretion of IFNγ and TNF results in transcriptional reprogramming of melanoma cells and induction of neural crest-like gene signatures (Grasso et al., 2021; Kim et al., 2021). The findings in our study suggest that baseline T cell infiltration levels (prior to the onset of immunotherapy) can regulate the transcriptional cell state of melanoma cells and induce LRP2 expression. It is likely that increased IFNγ signaling following immunotherapy further drives LRP2 in melanoma cells, although this remains to be examined. It also remains unknown if IFNγ regulates LRP2 expression in other cancers or inflammatory conditions.

Based on the established function of LRP2 as a rapidly internalizing and recycling receptor with more than 60 identified ligands, we speculate that melanoma LRP2 modulates the cellular uptake and extracellular concentration of its ligands, thereby impacting the composition of the tumor microenvironment. In addition, our previous work demonstrated endocytic functionality of melanoma LRP2 (Andersen et al., 2015). The observation that LRP2 correlates with T cell infiltration in melanoma raises the possibility that LRP2 function promotes T cell-mediated tumor immune responses. An interesting LRP2 ligand in this context is vitamin D-binding protein, which transports vitamin D (25 hydroxyvitamin D_3_ = 25(OH)D_3_) to target cells, thus facilitating the intracellular conversion of the precursor 25(OH)D_3_ to the active vitamin D (1,25(OH)_2_D_3_) (Nykjaer et al., 1999), as increased intratumoral vitamin D receptor signaling has recently been linked to T cell infiltration and immunotherapy response in melanoma (Giampazolias et al., 2024; Muralidhar et al., 2019). We set out to investigate the impact of LRP2 on the tumor microenvironment experimentally. However, despite extensive efforts, we were not able to identify Lrp2-expressing murine melanoma models for *in vivo* experiments. Examining the functional impact of LRP2 on the tumor microenvironment will therefore probably require the use of more advanced human *ex vivo* cancer models (Revach et al., 2024; Sun et al., 2025; Sun et al., 2023) in future studies.

Immunohistochemical analysis demonstrated higher LRP2 expression in stage I melanoma compared to stage II-or stage III melanoma. Although we observe decreasing LRP2 expression with disease progression in primary melanoma, we note that we previously demonstrated expression of *LRP2* in melanoma metastases (Andersen et al., 2015). In present study, we also observe high *LRP2* transcript levels in the melanoma metastases that compose the TCGA SKCM dataset. We speculate that LRP2 might be induced in melanoma metastases with a high degree of immune infiltration and IFNγ signaling.

In summary, the present study defines the molecular features of LRP2 expressing melanomas and identifies IFNγ signaling as a novel strong positive regulator of LRP2 in melanoma. Our findings encourage future studies on LRP2 in settings with increased IFNγ signaling, in particular in melanoma metastases following the onset of immunotherapy.

## Materials and Methods

### Patient samples

Patient samples (n=150) from a previously established cohort of Danish patients with primary cutaneous melanoma (n=385)(Bonnelykke-Behrndtz et al., 2017) were included for immunohistochemical analysis. Data on pathological parameters and follow-up information on the included patients were collected from electronic patient files. Clinical stage was revised according to the American Joint Committee on Cancer’s 8^th^ edition melanoma staging criteria. Patient samples were included in pairs matched according to ulceration, Breslow thickness, and age. Pairs were randomly selected from available sections in the previously established cohort. Clinicopathological characteristics of the included patients are presented in Supplementary Table 2. The Regional Committee for Health Research in the Central Denmark Region approved the study.

### PCA and subtype classification

Principal component analysis (PCA) was performed on log2 transformed gene expression values with an offset of 1. PCA was performed on mean centered data in R using prcomp. PCA plotting was performed using ggplot2 (Wickham, 2016). Cell line subtype classifications from the original publication were provided by Professor Thomas G. Graeber (University of California, Los Angeles, CA, USA) (Tsoi et al., 2018).

### RNA isolation, sequencing, data processing and subtype classification for FM3 and FM88

Total RNA was isolated from melanoma cells using the RNeasy Mini Kit (Qiagen). Bulk RNA sequencing library preparation (polyA-selected), quality control and paired-end 100 base pair sequencing (40 million paired reads per sample) on a DNBseq Sequencing system were performed at BGI Copenhagen. Paired sequencing files were processed to transcript level counts with kallisto (Bray et al., 2016) using ENSEMBL homo sapiens transcriptome v94 as the transcriptome index. Transcript level counts were summed to gene level counts using tximport, normalized using the trimmed mean of M-values (TMM) and transformed to log2 counts per million (CPM) using limma-voom (Law et al., 2014; Ritchie et al., 2015) in edgeR (Robinson et al., 2010). CPM values were used for subtype classification of melanoma cell lines FM3 and FM88 using a support vector machine “top-scoring pairs”-based approached (Tsoi et al., 2018) with scripts provided by Professor Thomas G. Graeber (University of California, Los Angeles, CA, USA). FM3 and FM88 were classified as transitory.

## Data availability

TCGA data was downloaded from the UCSC Xena data portal (https://xena.ucsc.edu). TCGA RNA sequencing data were processed using the Toil pipeline (Vivian et al., 2017), which uses STAR (Dobin et al., 2013) to generate alignments (reference genome GRCh38) and performs quantification using RSEM (Li & Dewey, 2011) and Kallisto (Bray et al., 2016). RNA sequencing data were downloaded as RSEM-based normalized gene quantifications and then log2-transformed with an offset of 1. For all analyses, the TCGA SKCM dataset was filtered to include only metastatic samples (n = 366). UCLA melanoma cell line RNA sequencing data were available as processed fragments per kilobase million (FPKM) values on GEO under accession number GSE80829 (Tsoi et al., 2018) and GSE154996 (Grasso et al., 2021). All data generated for this study are available from the authors upon reasonable request.

### Cell culture

Human melanoma cell lines were cultured under standard conditions in a humidified incubator with 5% CO_2_ at 37 °C. FM3 and FM88 were from Professor Per Guldberg (The Danish Cancer Society). SKMEL28, WM2664 and WM115 were from Professor Boe S. Sørensen (Aarhus University Hospital). A375 and RPMI7951 were purchased from American Type Culture Collection (ATCC). FM3, FM88, SKMEL28, WM2664 and WM115 were cultured in RPMI1640 (Sigma Aldrich) and A375 and RPMI7951 were cultured in DMEM (Gibco). Cell culture media was supplemented with 10% FBS (Gibco) and 100 units/mL penicillin and streptomycin (Sigma Aldrich).

### IFN**γ** stimulation

Cells were plated and cultured for 24 hours prior to IFNγ stimulation. Recombinant human IFNγ (PeproTech) was used at 50 ng/mL for 24-hour time course and 48-hour stimulation experiments and at 25 ng/mL for 5-week time course experiments. Cell culture medium was replaced with fresh medium containing IFNγ every 2-3 days for the 5-week time course experiments.

### RT-qPCR

Cells were washed in PBS and lysed with RLT buffer (Qiagen). RNA was purified using the RNeasy Mini Kit (Qiagen) following the manufacturers protocol with on-column DNase digestion and eluted in nuclease-free water. Purified RNA concentration was measured on a Nanodrop 100 Spectrophotometer (Thermo Fisher Scientific). For all samples, a total of 1440 ng RNA in a 20 μl reaction volume was used for cDNA synthesis. cDNA synthesis mixtures were prepared using the High Capacity RNA-to-cDNA kit (Applied Biosystems) and cDNA synthesis (37 °C for 1 hour) and denaturation (95 °C for 5 minutes) steps were run on a Verity Thermal Cycler (Applied Biosystems). RT-qPCR was performed using TaqMan^TM^ Fast Advanced Master Mix and assays (Life Technologies) according to the manufacturers protocol. RT-qPCR was run on a QuantStudio Real-Time PCR system (Thermo Fisher Scientific). Cycle threshold (Ct) values were determined using a baseline-subtracted normalized reporter value (ΔRn) of 0.4 for all genes. Ct values above 36 were considered as not detected. ΔCt values were calculated as the difference between the geometric mean of Ct values between the gene of interest and the reference gene *CASC3* (Christensen et al., 2020). Relative quantification (RQ) values were calculated using the formula 2^-ΔΔCt^. Ct values for *GAPDH* measured on a stock preparation of cDNA from FM88 was used as an interplate control. Taqman^TM^ probes used in this study: *CASC3* (Hs00201226_m1), *MITF* (Hs01117294_m1), *MLANA* (Hs00194133_m1), *LRP2* (Hs00189742_m1), *NGFR* (Hs00609976_m1), *AXL* (Hs01064444_m1) and *GAPDH* (Hs02758991_g1).

### Immunoblotting

Cell pellets were lysed in “membrane buffer” (20 mM CaCl_2_, 10 mM MgCl_2_, 100 mM HEPES, 1400 mM NaCl, pH 7.8) with 1% Triton X-100 (Merck) and cOmplete^™^ Mini Protease Inhibitor Cocktail for 15 minutes on ice. Cell lysates were centrifuged at 13.000 rpm for 15 minutes at 4 °C and supernatant was collected. Total protein concentration was determined using the Pierce^™^ BCA Protein Assay kit (Thermo Scientific). 5% SDS sample buffer was added to samples which were then boiled for 5 minutes. Protein samples were loaded onto non-reducing NuPAGE^™^ 3-8% Tris-Acetate gels (Invitrogen) along with SeeBlue^™^ Plus Pre-stained Protein Standard and run at 200 V for 75 minutes. Protein blotting onto a polyvinylidene difluoride (PVDF) membrane was performed overnight at 4 °C using a constant current of 100 mA in a Mini Tank Transfer Unit (GE Healthcare Life Sciences). Following blotting, PVDF membranes were blocked in 1x Tris-buffered saline with 2% Tween® 20 (Sigma Aldrich) (TBST) overnight at 4 °C. Primary antibodies were diluted in 5 % w/v nonfat dry milk, and incubation was performed for two hours at room temperature. Following washing, incubation with horseradish peroxidase (HRP)-conjugated secondary antibody (SIGMA) was performed for one hour at room temperature. Pierce ECL Western Blotting Substrate (Thermo Scientific) was added and protein bands were visualized on a LAS-3000 Imaging System (FujiFilm). Primary antibodies were mouse monoclonal anti-human megalin (Rasmussen et al., 2023), mouse monoclonal anti-clathrin (gift from Linton M. Traub) (Nathke et al., 1992) and anti-β actin (A5441, Sigma).

### Gene Set Enrichment Analysis (GSEA)

Genes were ranked based on their correlation to LRP2 in TCGA SKCM metastases. Gene set enrichment analysis (GSEA) was performed on the ranked gene list using the fgsea package (https://www.biorxiv.org/content/10.1101/060012v3) in R. Gene sets included HALLMARKS and GO Biological Process (BP). Enrichment plots were based on running enrichment scores extracted from the ClusterProfiler (Yu et al., 2012) package in R.

### MCPcounter and T cell inflammation scores

Microenvironment cell population counter (MCPcounter) was applied to the TCGA SKCM gene expression matrix to calculate absolute abundance scores for eight major immune cell types (T cells, CD8 T cells, cytotoxic lymphocytes, NK cells, B cells, monocytic lineage cells, myeloid dendritic cells and neutrophils) as well as fibroblasts and endothelial cells (Becht et al., 2016). MCPcounter absolute abundance scores were mean-centered (z-score) for each cell type and compared between *LRP2*-high and *LRP2*-low groups (upper quartile cutoff). T cell inflammation scores for TCGA SKCM were calculated as aggregated mean-centered scores (z scores) of a previously defined T cell inflammation gene signature: *CD8A*, *CCL2*, *CCL3*, *CCL4*, *CXCL9*, *CXCL10*, *ICOS*, *GZMK*, *IRF1*, *HLA-DMA*, *HLA-DMB*, *HLA-DOA* and *HLA-DOB* (Spranger et al., 2015). T cell inflammation-low and T cell inflammation-high stratification was based on a median cutoff.

### Four-color multiplexed immunohistochemistry and digital image analysis

Formalin-fixed paraffin-embedded (FFPE) melanoma tumors were sectioned and stained at the Department of Pathology at Aarhus University Hospital. FFPE samples were sectioned at a thickness of 3 μM and mounted on TOMO® Adhesion Microscope Slides (Matsunami Glass Ind., Ltd.). Slides were dried at 60 °C for one hour prior to staining. Sequential staining was performed on a DISCOVERY ULTRA (Roche) in the following order: CD8 (clone C8/144B, Dako, 1:150) detected with DISCOVERY ChromoMap DAB kit (RUO)(Roche), CD4 (clone SP35, Ventana, ready to use) detected with DISCOVERY Purple HRP (Roche), LRP2 (protein G-purified polyclonal rabbit anti-human LRP2 antibody (Andersen et al., 2015) (1:100) detected with DISCOVERY Teal HRP (Roche), and SOX10 (clone SP267, Ventana, ready to use) detected with DISCOVERY Yellow AP (Roche). Upon staining, tissue sections were scanned at 20x magnification on a NanoZoomer S60 Digital Slide Scanner (Hamamatsu Phototonics).

Image analysis was performed using VisioPharm software (Visiopharm A/S). We applied a threshold-based color classification to label positive staining for the four markers: CD8 (DAB), CD4 (purple), LRP2 (teal) and SOX10 (yellow). First, a global tumor region of interest (ROI) was outlined manually, which included bulk tumor and surrounding lymphocytes, while excluding epidermis, large blood vessels, heavily pigmented regions, and artifacts from sample preparation. Second, an automated image analysis pipeline was applied, which calculated the area fraction of CD8 and CD4 staining within the global tumor ROI, used the SOX10 label to define a SOX10+ melanoma cell ROI (with manual correction if needed), and calculated the area fraction of LRP2 staining within the SOX10+ melanoma cell ROI. Following quality control of sectioning and staining of the 150 sections, there were 141 sections available for measurement of LRP2 (SOX10+ area fraction) and 132 sections available for measurement of both LRP2 (SOX10+ area fraction) and T cells (CD4 and CD8 tumor area fraction). T cell inflammation stratification was based on aggregated CD4 (tumor area fraction) and CD8 (tumor area fraction) scores using a median cutoff. Two sections were excluded from the assessment of LRP2 expression and clinicopathological parameters due to missing information, leaving 139 sections available for these analyses.

### Statistical analysis and data visualization

Data handling, statistical analysis and data visualization was performed in and R version 4.0.4 and Prism version 9.2.0. Digital image analysis of multiplex immunohistochemistry was in VisioPharm.

## Figure Legends

**Supplementary Figure 1.**
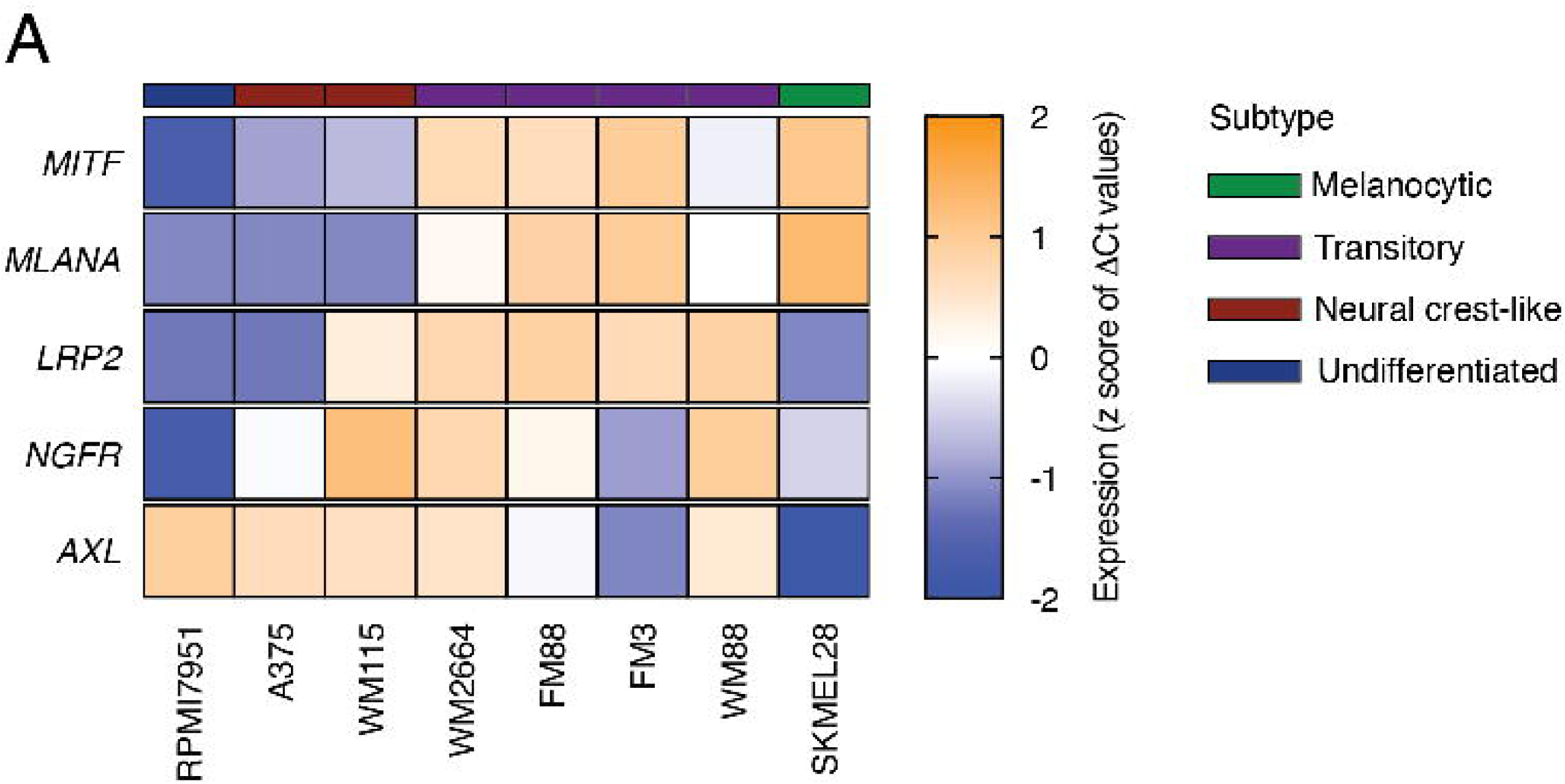
Expression of melanoma differentiation cell state marker genes in melanoma cell lines. **A.** Heatmap of melanoma differentiation cell state marker gene expression determined by RT-qPCR across the indicated melanoma cell lines. Z scores for ΔCt values normalized to *CASC3* are shown.

**Supplementary Figure 2.**
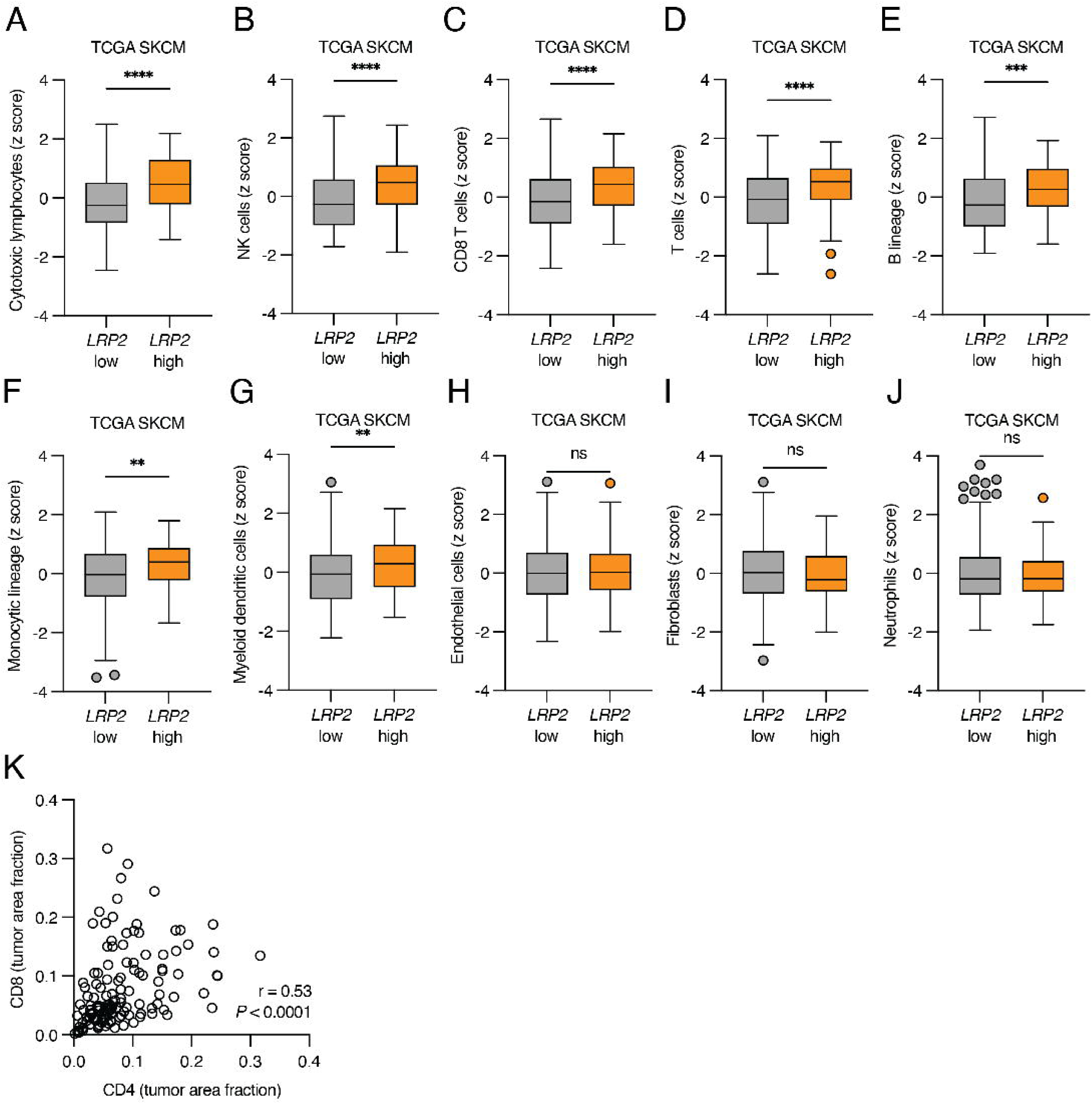
LRP2-expressing melanomas have increased immune infiltration. **A-J.** Boxplots of MCPcounter cell type scores in *LRP2*-low and *LRP2*-high (upper quartile cutoff) TCGA SKCM (unpaired t-test). The boxplot lines represent the lower quartile, median, and upper quartile. Whiskers extend to 1.5 times above or below the interquartile range. **K.** Scatter dot plot of CD4 and CD8 (tumor area fraction) values from multiplex immunohistochemistry of primary melanoma tumors. Statistics for Spearman’s correlation are shown. **P < 0.01. ***P < 0.001. ****P < 0.0001. ns = not significant.

**Supplementary Figure 3.**
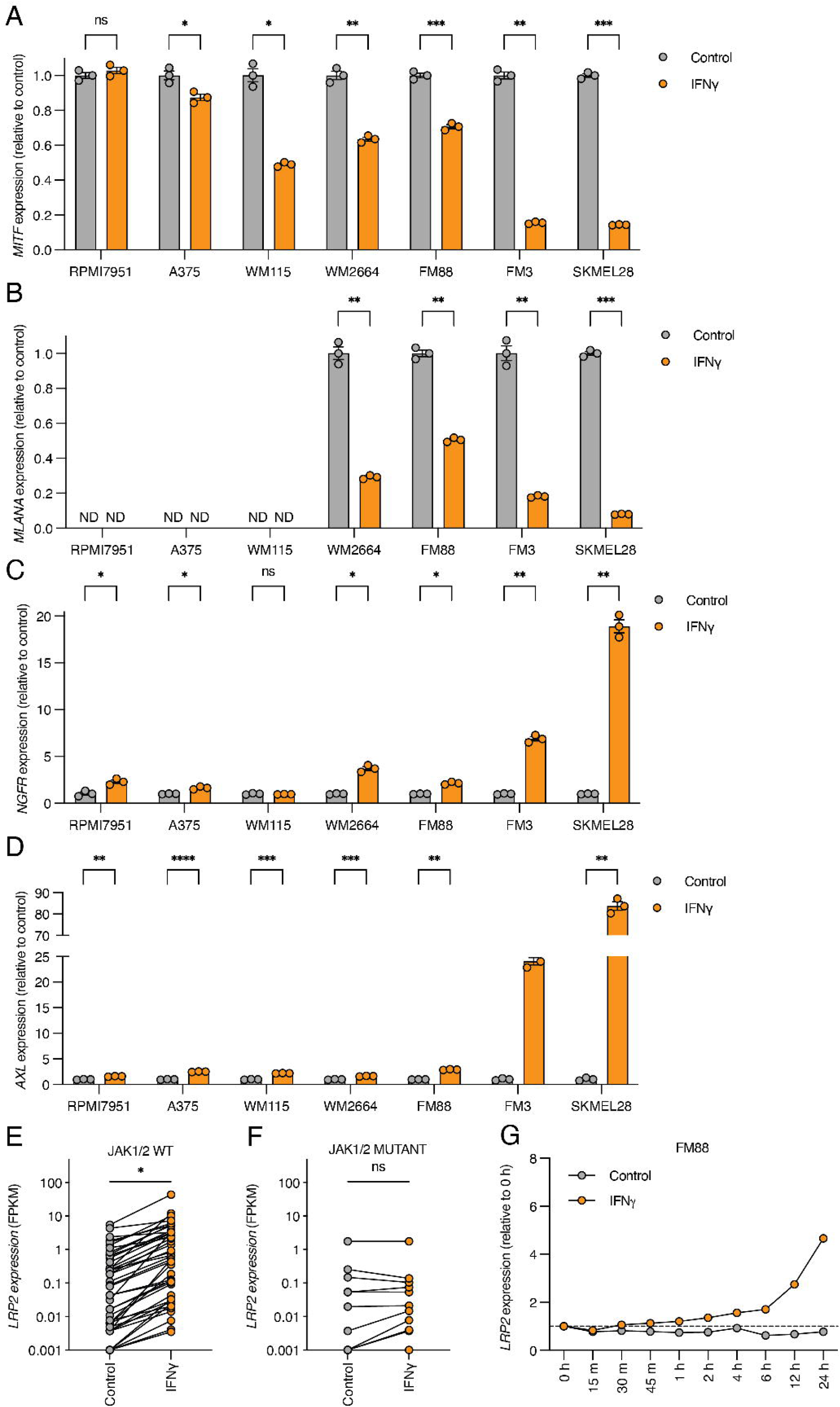
IFN**γ** increases LRP2 expression in melanoma cells. **A-D.** Bar plot of gene expression (relative quantities) determined by RT-qPCR in the indicated melanoma cell lines treated with vehicle control or IFNγ for 48 hours (multiple unpaired t-tests). Mean (bars) ± SEM (error bars) and individual values (circles) are shown (n = 3 technical replicates). **E-F.** *LRP2* expression (FPKM) determined by bulk RNA-sequencing in JAK1/2 WT (E) or JAK1/2 mutant (F) melanoma cell lines treated with vehicle control or IFNγ (paired t-test). **G.** *LRP2* expression (relative quantities) determined by RT-qPCR during a 24-hour time course in cells treated with vehicle control or IFNγ. Mean (circles) ± SEM (error bars) are shown (n = 3 technical replicates). No error bars are displayed when the error is smaller than the size of the bar. *P < 0.05, **P < 0.01, ***P < 0.001, ****P < 0.0001. ND = not detected. ns = not significant.

**Supplementary Figure 4.**
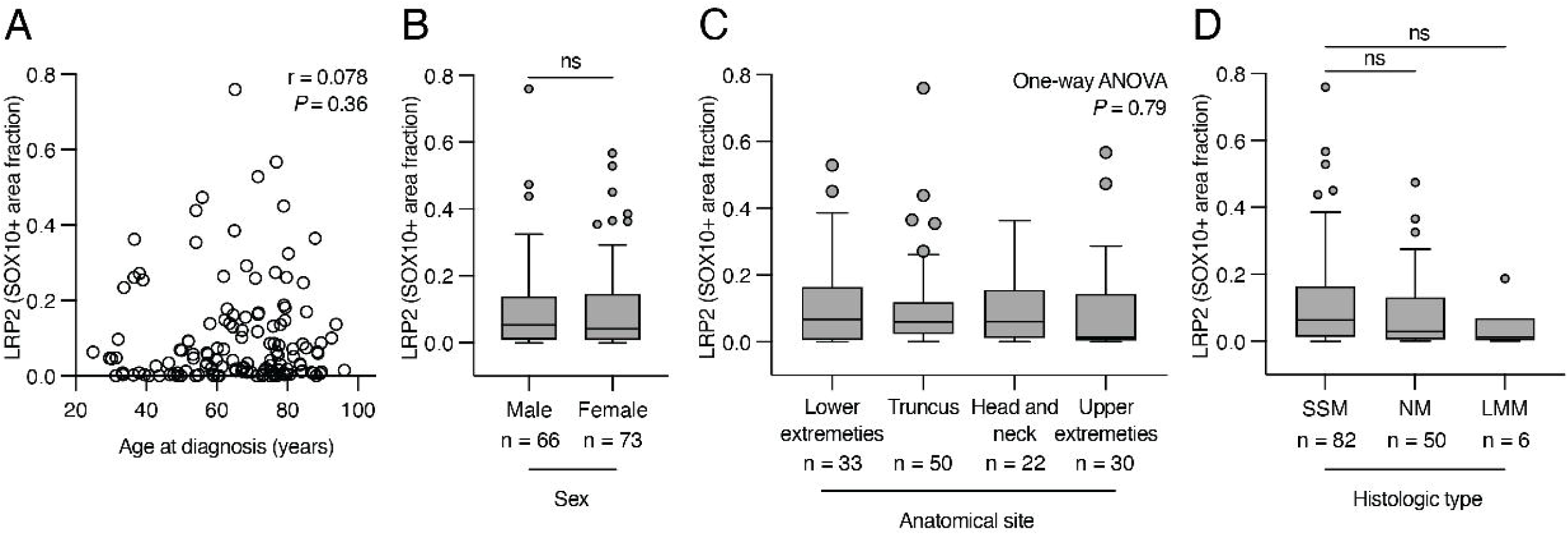
LRP2 expression is not associated with patient age, sex, anatomical site or histological type in primary melanoma. **A.** Scatter plot of patient age at diagnosis (x axis) and LRP2 (SOX10+ area fraction) expression (y axis) in primary melanoma. Statistics for Spearman’s correlation are shown. **B.** Boxplot of LRP2 (SOX10+ area fraction) expression by patient sex in primary melanoma (Mann-Whitney test). **C.** Boxplot of LRP2 (SOX10+ area fraction) expression by anatomical site in primary melanoma (one-way ANOVA). **D.** Boxplot of LRP2 (SOX10+ area fraction) by histological type in primary melanoma (Mann-Whitney test). The boxplot lines represent the lower quartile, median, and upper quartile. Whiskers extend to 1.5 times above or below the interquartile range. ns = not significant. SSM = superficially spreading melanoma. NM = nodular melanoma. LMM = lentigo maligna melanoma.

## Acknowledgements

This work was supported by grants from The Danish Cancer Society (M.Q.R), The Danish Cancer Foundation (M.Q.R), Graduate School of Health at Aarhus University (M.Q.R), The Novo Nordisk Foundation (M.M), The Eva and Henry Fraenkel Memory Foundation (M.M), The Dagmar Marshall Foundation (M.M), The A.P. Moller Foundation (M.M), The Einar Willumsen Foundation (M.M), Architect Holger Hjortenberg Foundation (M.M), The Max Woerzner and Inger Woerzner Foundation (M.M). The funding bodies had no role in the design of the study, and collection, analysis, and interpretation of the data, or in writing the manuscript. The authors thank Kristina Lystlund Lauridsen for discussions on digital pathology and also thank all members of the Mette Madsen laboratory at Aarhus University.

