## Supplementary Table 1 for "LRP2 expression in melanoma is associated with a transitory cell state, increased T cell infiltration, and is upregulated by IFNγ signaling"

**Supplementary Table 1:** Gene set enrichment analysis of *LRP2* correlated genes in TCGA SKCM. HALLMARK and GO BP gene sets were analyzed. Gene sets are ranked by NES in decreasing order. The 10 most upregulated and 10 most downregulated pathways (based on NES) are shown.

| Gene set | Gene set  size | Enrichment score | Normalized enrichment score | P value | P value adjusted | -log10(P value adjusted) |
| --- | --- | --- | --- | --- | --- | --- |
| HALLMARK INTERFERON GAMMA RESPONSE | 198 | 0,72 | 2,81 | 8,79E-43 | 4,03E-40 | 39,40 |
| HALLMARK INTERFERON ALPHA RESPONSE | 95 | 0,75 | 2,78 | 2,94E-25 | 3,03E-23 | 22,52 |
| GOBP ADAPTIVE IMMUNE RESPONSE | 430 | 0,68 | 2,76 | 2,96E-72 | 1,22E-68 | 67,91 |
| GOBP LYMPHOCYTE MEDIATED IMMUNITY | 280 | 0,68 | 2,73 | 5,92E-49 | 4,88E-46 | 45,31 |
| GOBP B CELL MEDIATED IMMUNITY | 138 | 0,71 | 2,72 | 1,97E-29 | 2,90E-27 | 26,54 |
| HALLMARK ALLOGRAFT REJECTION | 195 | 0,68 | 2,69 | 2,42E-35 | 5,54E-33 | 32,26 |
| GOBP POSITIVE REGULATION OF T CELL PROLIFERATION | 100 | 0,72 | 2,67 | 5,73E-22 | 4,73E-20 | 19,33 |
| GOBP REGULATION OF LYMPHOCYTE MEDIATED IMMUNITY | 165 | 0,69 | 2,67 | 4,48E-31 | 6,84E-29 | 28,17 |
| GOBP INTERFERON GAMMA PRODUCTION | 113 | 0,71 | 2,66 | 2,90E-23 | 2,72E-21 | 20,57 |
| GOBP POSITIVE REGULATION OF CELL KILLING | 62 | 0,76 | 2,66 | 8,98E-18 | 5,00E-16 | 15,30 |
| … | … | … | … | … | … | … |
| GOBP RIBOSOMAL SMALL SUBUNIT BIOGENESIS | 67 | -0,52 | -2,38 | 3,49E-08 | 6,60E-07 | 6,18 |
| GOBP RIBOSOME ASSEMBLY | 57 | -0,56 | -2,52 | 6,59E-08 | 1,19E-06 | 5,93 |
| HALLMARK E2F TARGETS | 195 | -0,46 | -2,53 | 3,16E-16 | 1,53E-14 | 13,81 |
| GOBP RIBOSOMAL LARGE SUBUNIT ASSEMBLY | 25 | -0,71 | -2,60 | 2,14E-07 | 3,44E-06 | 5,46 |
| GOBP CYTOPLASMIC TRANSLATION | 144 | -0,51 | -2,63 | 1,84E-15 | 8,24E-14 | 13,08 |
| GOBP RRNA METABOLIC PROCESS | 239 | -0,49 | -2,72 | 1,01E-22 | 9,29E-21 | 20,03 |
| HALLMARK MYC TARGETS V2 | 58 | -0,63 | -2,81 | 1,27E-11 | 3,83E-10 | 9,42 |
| GOBP RIBOSOMAL LARGE SUBUNIT BIOGENESIS | 68 | -0,62 | -2,82 | 3,08E-12 | 1,02E-10 | 9,99 |
| HALLMARK MYC TARGETS V1 | 194 | -0,51 | -2,82 | 1,30E-21 | 1,03E-19 | 18,99 |
| GOBP RIBOSOME BIOGENESIS | 280 | -0,50 | -2,90 | 7,96E-29 | 1,09E-26 | 25,96 |
