## Supplementary Table 2 for "LRP2 expression in melanoma is associated with a transitory cell state, increased T cell infiltration, and is upregulated by IFNγ signaling"

| **Supplementary Table 2:** Clinicopathological characteristics of the included patients. | | |
| --- | --- | --- |
| **Variables** | **Numbers (%)** | **Median (min. - max.)** |
| **Age at the time of diagnosis** |  | 67.7 (18.0 - 96.1) |
| **Sex** |  |  |
| Male | 66 (47) |  |
| Female | 73 (53) |  |
| **Location** |  |  |
| Head and neck | 22 (16) |  |
| Truncus | 50 (36) |  |
| Upper extremity | 30 (21) |  |
| Low extremity | 33 (24) |  |
| Unknown | 4 (3) |  |
| **Histological subtype** |  |  |
| Superficially spreading melanoma | 82 (59) |  |
| Nodular melanoma | 50 (36) |  |
| Lentigo meligna melanoma | 6 (4) |  |
| Unknown | 1 (1) |  |
| **Breslow thickness** |  |  |
| <1.0 mm | 11 (8) |  |
| 1.1 - 2.0 mm | 30 (22) |  |
| 2.1 - 4.0 mm | 73 (53) |  |
| >4.0 mm | 23 (16) |  |
| Unknown | 2 (1) |  |
| **Ulceration** |  |  |
| Yes | 64 (46) |  |
| No | 74 (53) |  |
| Unknown | 1 (1) |  |
| **Clinical Stage, AJCC*** 8th edition** |  |  |
| I | 26 (19) |  |
| II | 68 (49) |  |
| III | 39 (28) |  |
| IV | 4 (3) |  |
| Unknown | 2 (1) |  |
| ***American Joint Committee on Cancer | | |
